# Two forms, two functions: functional strategies of parasitoid bristle flies and their larvae

**DOI:** 10.1101/2025.09.08.674822

**Authors:** Moreno Di Marco, Michela Gabrieli, Lara Marcolin, Noemi Di Lorenzo, Pierfilippo Cerretti, Luca Santini

## Abstract

Species with complete metamorphosis undergo substantial changes in ecological functions, especially parasitoid insects with parasitic larvae and free-living adults. While parasitoids play essential ecosystem roles, limited knowledge is available about their shifting functional strategies. Here we focus on parasitoid bristle flies (Diptera: Tachinidae) to investigate the relationship between larval *vs* adult functional strategies. We retrieved trait data for 767 European species and defined functional trait spaces for larvae and adults. We then measured both functional distinctiveness (rarity in functional trait combination) and specialisation (variety of resources consumed). We found little correspondence in the functional distinctiveness of adults and larvae, with highly distinct larvae generating either functionally distinct or functionally common adults. In contrast, only specialised larvae (attacking a limited number of hosts) give origin to specialised adults (feeding on a limited number of flowers). This suggests selective pressure towards specialisation might act synergistically across life stages, if trophic resources are restricted in space for both the larva (e.g. caterpillar host) and adult (e.g. flowers). Global change can generate complex patterns of functional homogenisation in parasitoids, which can occur at different (or both) life stages and lead to ecosystem-wide consequences: from the outbreak of herbivore insects to the loss of pollination capacity.

## Main

Functional diversity is a key property of biological communities which captures the variation in species’ ecological roles. From a theoretical perspective, functional diversity represents “what species do” ^1^, although in practice ecological roles are typically inferred from functional traits. These traits – which encompass aspects of ecology, physiology, and life history – reflect species adaptations to the environment and are linked to individual fitness and overall species persistence within a community ^2, 3^. High functional trait diversity within a biological community reflects the presence of species that exploit diverse resources and play complementary roles ^4^. This is often compared to taxonomic diversity or phylogenetic diversity ^5^, which respectively quantify the number of taxa in a community and the amount of evolutionary history they encompass, estimated from phylogenetic trees. Functional diversity is a multi-dimensional concept which encompasses several aspects of a species’ roles in an ecosystem, it is a complex multi-faceted property of biological communities ^6^.

Species contributions to functional diversity can be described through multiple metrics, such as functional specialisation, which represents the species’ distance from the “intermediate” condition in the trait space, or functional distinctiveness, which represents the difference between a given species and all other species in the community ^7^. These dimensions can offer complementary information and be decoupled: for example, highly specialised species determine the limits of the functional space of a community while highly distinct species may occupy underrepresented portions of functional space ^7, 8^. But the situation complicates even further for species undergoing metamorphosis, which shifts their functional roles across life stages. In this case, the two stages of the same species might have different functional strategies (e.g. specialist *vs* generalist) not just different functional roles. Parasitoid insects, such as many wasps (Hymenoptera) and flies (Diptera), are a particular example of organisms living a dual life, undergoing complete functional shifts across their life. These species shift from parasitic larvae – feeding on a wide range of invertebrate hosts, such as arthropods, molluscs, and annelids – to free-living adults. During their larval stage, parasitoids play a key role in regulating host population dynamics ^9^, often acting as biological control agents ^10, 11^. As adults, they typically feed on floral nectar and/or other carbohydrate-rich substances, delivering important contributions to the pollination networks of several ecosystems.

Bristle flies (Diptera: Tachinidae) are a megadiverse group of parasitoid insects that primarily exploit herbivorous insects as hosts, especially caterpillars^12, 13^. Adult bristle flies play important roles as pollinators, especially in montane ecosystems where they may complement, or even replace, other pollinators such as bees ^14^. Yet, bristle flies are particularly sensitive to environmental changes ^15^, due to both direct and indirect environmental impact mediated by changes in their hosts’ dynamics. Recent work demonstrated that global change has altered the functional diversity of bristle fly communities across Europe ^16^, flattening the elevational gradient of larval diet specialisation that characterised these species in the 1960s. However, this assessment was limited to the larval diet spectrum, disregarding adult traits and other dimensions of functional diversity. It is indeed possible that species exhibit different levels of specialisation across their two life stages, and/ or different levels of distinctiveness.

Here we explore the dual life of parasitoid bristle flies from a functional perspective, investigating multiple dimensions of functional strategies across the larval and adult stages. We retrieved functional trait data for 767 species of European bristle flies, out of approximately 900 ^17, 18^, covering all tribes and nearly all genera currently recorded from the European fauna. We then measured the functional distinctiveness and functional specialisation of each species at both larval and adult stages. For simplicity, we attributed to the “larval” stage all traits connected with the species’ development strategy, including those related to the egg. First, we test whether there is a tradeoff between the functional distinctiveness of larvae and adult stages. Second, we repeat this test focusing only on traits related to diet. Third, we focus on diet specialisation by looking at bristle flies that parasitise lepidopterans, to test whether functional specialisation in one life stage constrains the degree of specialisation in the other. Our aim is to identify correspondences of functional strategies between the larval and adult life stages of each species, as well as correlations between functional distinctiveness and functional specialisation. This integrative approach provides a novel framework to understand how multi-stage organisms, with complex life cycles contribute to functional diversity and how functional homogenisation may unfold differently across life stages.

## Results

### Functional distinctiveness in larvae *vs* adults

We found the functional morphospace of European bristle fly species to be patchy, with several “high-density” areas (i.e. areas including many species sharing common strategies), but also gaps in between indicating non-functional combinations of traits (Fig. 1). This pattern is reflected at both the larval and adult stages, and several species occupy areas of high functional density at both life stages. For example, *Pales pavida* – a generalist species able to colonise multiple environments, feeding on a variety of hosts as a larva, and showing generic feeding strategy as an adult – occupies high-density portions of the functional space at both life stages. This indicates that its functional traits are generally shared by several other species, both as a larva and as an adult. Instead, *Siphona collini –* whose larvae specialise in attacking a single family of lepidopterans (Noctuidae), and whose adults possess highly elongated and geniculate mouthparts specialised for feeding on flowers with a deep corolla (Fig. 1D) – occupies a high-density portion of the larval functional space but a low-density portion of the adult functional space. These examples reflect a shift from a relatively common trait combination at the larval stage to functionally uncommon adult traits, which is more in general reflected in a lack of correspondence between larval vs adult positioning in the morphospace (Extended Data Fig. 1).

**Figure 1.**
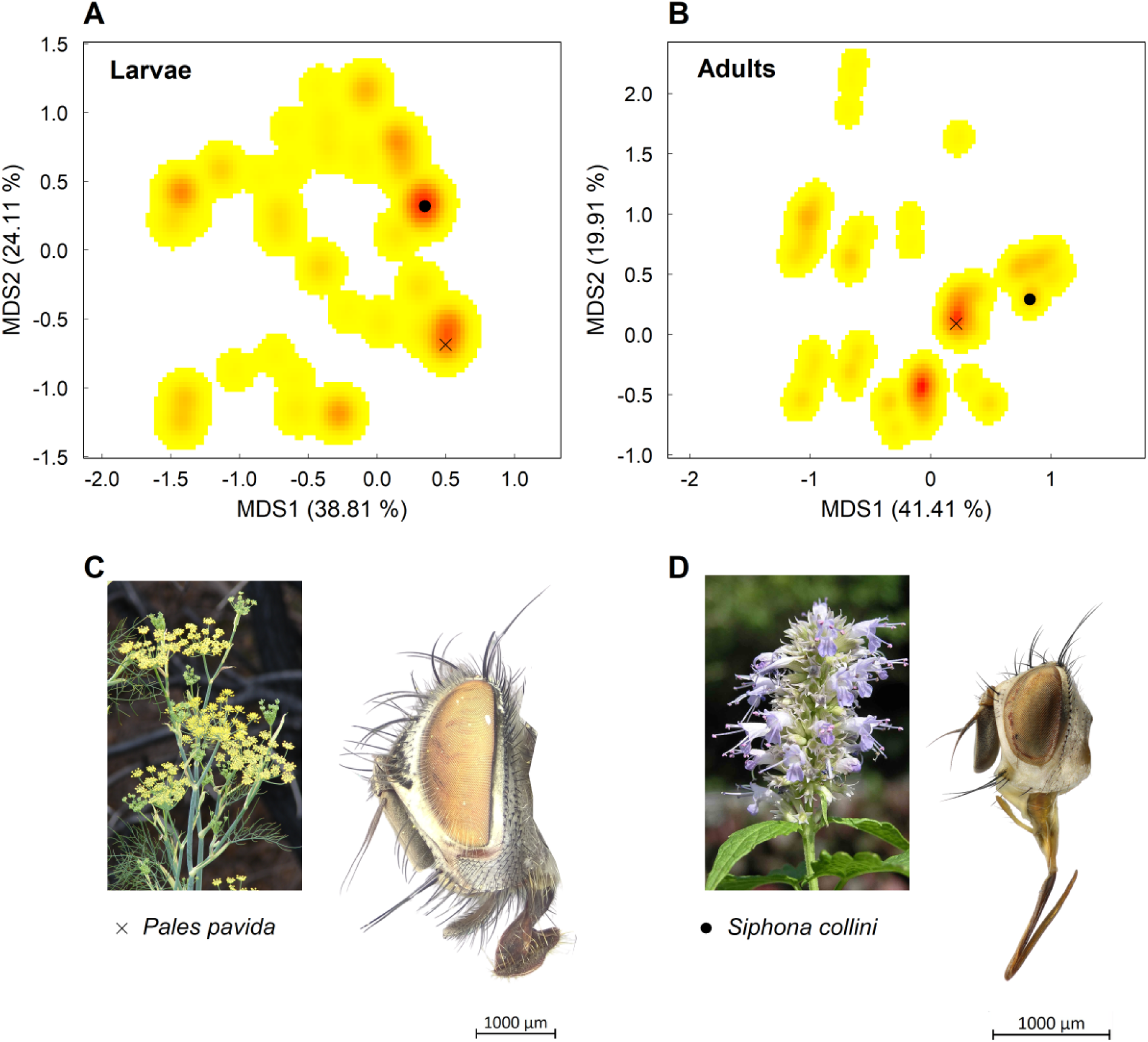
Two-dimensional density heatmaps of the functional morphospace for European bristle fly as (A) larvae and (B) adults, based on the first two axes of a non-metric multidimensional scaling (MDS) ordination. Warmer colors indicate areas with higher species density. Two representative species are highlighted: (C) *Pales pavida*, whose adults are characterized by broad mouthparts which indicate generalist nectar feeding (e.g. on *Foeniculum vulgare*, represented in figure); and (D) *Siphona collini*, with elongated, geniculate mouthparts typical of specialised species feeding on deep corollae (e.g. *Mentha spicata*, represented in the figure). Flower images retrieved from www.flickr.com. Photo of *S. collini* and copyright by Göran Liljeberg.

Similar to what we found when looking at commonness in the morphospace, we found that the functional distinctiveness of bristle fly species at larval and adult stages was largely uncorrelated (Fig. 2A). In particular, we found that a high degree of distinctiveness of parasitic bristle fly larvae does not necessarily correspond to a high degree of distinctiveness in free-living adults, and vice versa, even if we did find that only highly distinct larvae give origin to highly distinct adults. In contrast, when only looking at diet-related functional traits (i.e. attacked hosts for the larvae and mouthparts for the adults), the relationship between larval and adult functional distinctiveness showed a triangular pattern (Fig. 2B). In this case, no species displayed highly distinct larvae and highly distinct adults, i.e. high levels of distinctiveness are only achieved at one of the two life stages. Similar patterns were found when looking only at the subset of lepidopteran-feeder bristle flies (Extended Data Fig. 2), for which detailed host information (i.e. comprehensive list and characteristics of the host species) was available.

**Figure 2.**
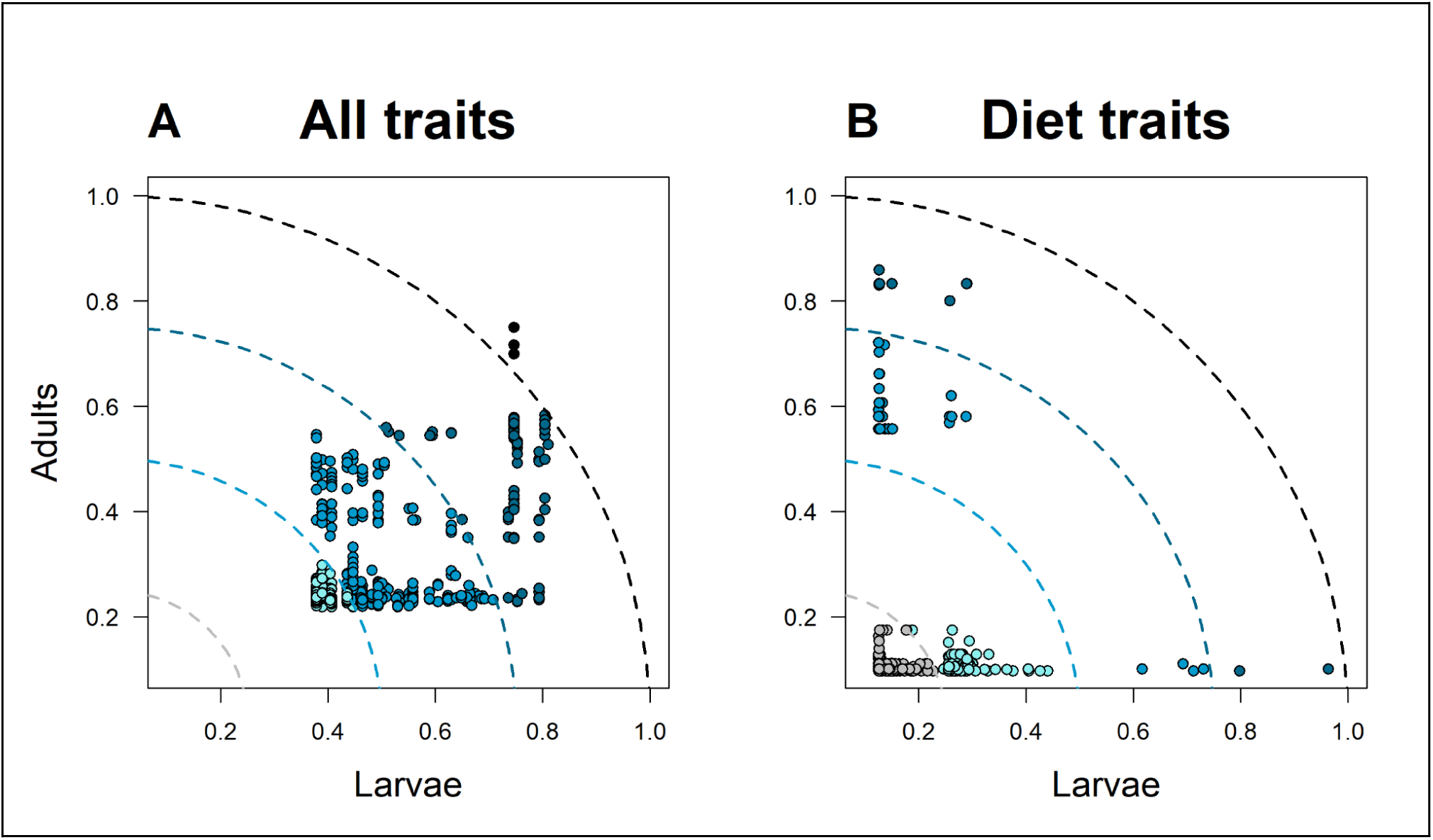
Functional distinctiveness of European bristle fly species at larval (x axis) *vs* adult (y axis) stage. Panels show results based on (A) all functional traits, and (B) only diet-related traits. The dashed lines represent increased distance from the origin (i.e. increased levels of distinctiveness).

### Functional specialisation in larvae *vs* adults

Distinctiveness indicates functionally rare combinations of traits, but does not necessarily represent ecological specialisation. We assessed diet specialisation for bristle fly larvae by looking at the defence strategies of caterpillar hosts. We focused on a subset of 302 bristle fly species parasitising lepidopteran hosts (caterpillars), hereafter “lepidopteran feeders”, for which detailed feeding information is available. We defined larval diet specialisation in terms of their ability to exploit “defence strategies” of the caterpillar hosts. Host defence strategies were assessed using traits associated with the following defensive mechanisms: aposematism, hairiness, gregariousness, and non-exposure (i.e. whether caterpillars forage exposed or remain concealed, either within vegetation structures or self-constructed cases). We assumed that specialists would be able to attack only a limited set of host defence strategies, whilst generalists would be able to exploit a wider range of strategies. In parallel, we assessed adult diet specialisation based on labellar shape, classified as broad and fleshy (generalist feeders; Fig. 1C), narrow, or elongated/geniculated, with labella folded against prementum (specialist feeders; Fig. 1D). We found that the vast majority of adult bristle flies are generalist feeders, while only few species show morphological adaptations consistent with floral specialisation, i.e. elongated/geniculated mouthparts (Extended Data Fig. 3A). At the larval stage most bristle flies attack caterpillars lacking any defensive traits (Extended Data Fig. 3B), which is consistent with the fact that those are the most common caterpillars in our dataset (Extended Data Fig. 3C). When looking at larval specialisation, measured as the number of defence strategies that a larva can exploit, we found that species span the full spectrum of specialism-generalism being able to exploit anywhere from 1 to 11 different defence strategies (Fig. 3). Overall, we found only 124 species specialise on a single defence strategy while all other species can exploit two or more strategies. There was no association between the level of larval specialisation (number of exploited defence strategies) and the type of defence strategy (Fig. 3), as virtually all strategies were attacked by both specialised bristle flies (those attacking one or two defence strategies) and generalist ones (those attacking many defence strategies). However, only specialised species (those attacking a single defence strategy) had a preference for the “no trait” strategy, i.e. represented by caterpillars with no defence mechanisms among those considered here (Fisher’s exact test p<<0.01; Extended Data Fig.s 4, 5).

**Figure 3.**
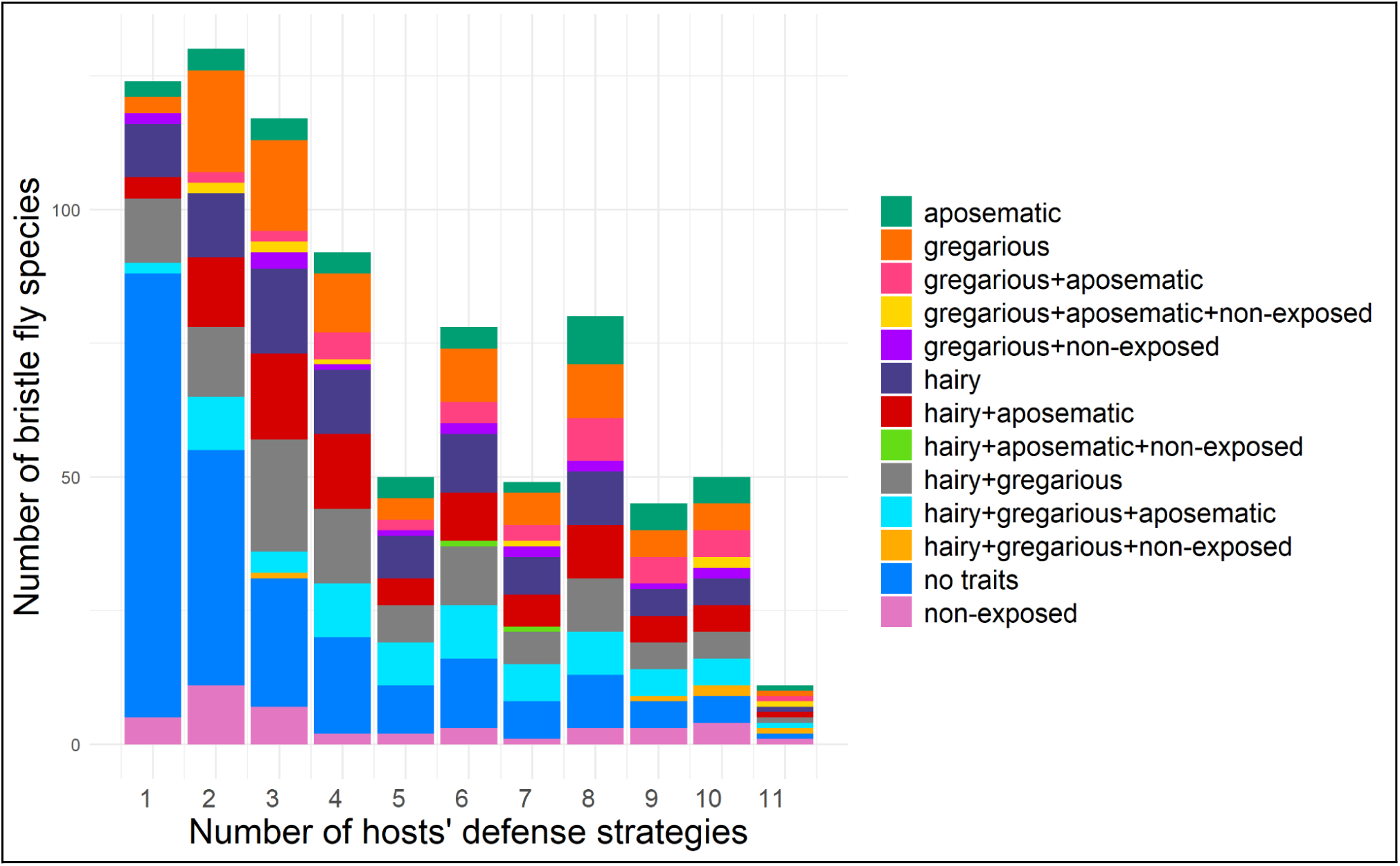
Number of bristle fly species that are able to exploit (as larvae) each number of lepidopteran defence strategies. This variable is used as a proxy for larval functional specialisation, with specialists only able to exploit a few defence strategies and generalists able to exploit many strategies.

When comparing diet specialisation in larval *vs* adult bristle flies, we found a triangular relationship (Fig. 4), horizontally specular to what we found with diet functional distinctiveness (Fig. 2B). While adult generalists could derive from larvae with any level of diet specialisation (low to high), specialised adults (those with elongated mouthparts) only derived from specialised larvae (those attacking only one or two strategies). We verified that our result on larval specialisation ￼￼was not an artefact of adults targeting a certain host for oviposition, as we repeated our analysis only on bristle fly species with indirect oviposition (i.e. adults not actively searching for a host) and found the same patterns (Extended Data Fig. 6). We also found the same patterns when discarding species that only predate on caterpillars with no defence mechanisms (Extended Data Fig. 7).

**Figure 4.**
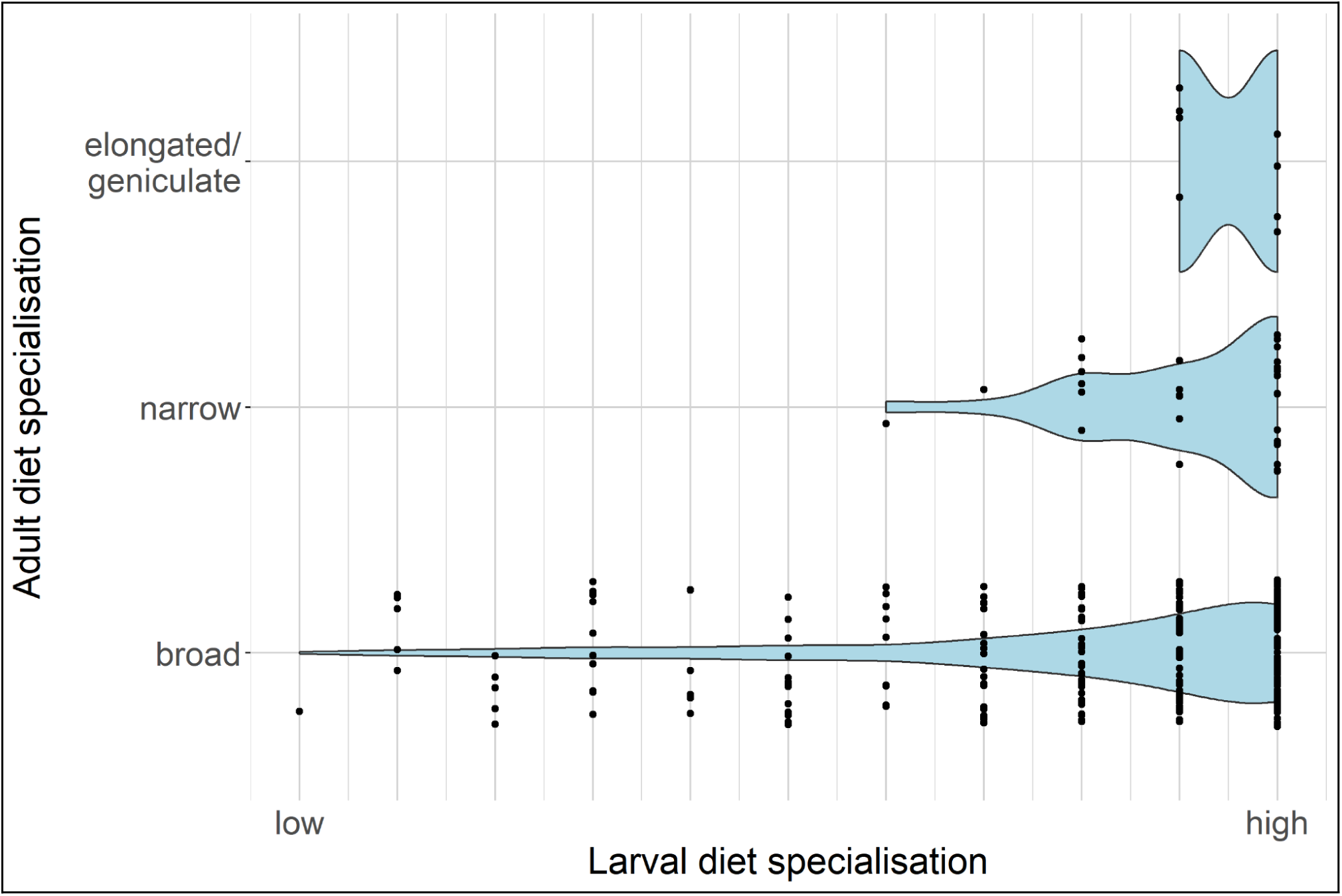
Relationship between diet functional specialisation at larval and adult stages. Larval specialisation is represented in terms of the number of hosts’ defence strategies attacked by the larva, in decreasing order (i.e. more specialised larvae can only attack a few host types). Adult specialisation is represented in terms of the shape of their mouth parts with “broad” being the most generalist form and “elongated” being the most specialised one.

## Discussion

Our analysis of stage-specific functional diversity for parasitoid bristle flies unveiled the decoupled relationship between adult and larval functional strategies. Although larval and adult functional distinctiveness were largely uncorrelated in our dataset, both diet distinctiveness and diet specialisation were not randomly distributed between the two stages of European bristle fly species. Specifically, we found that high levels of diet functional distinctiveness can only be achieved at either larval or adult stages, but not both. Instead, high levels of adult diet specialisation seem to correspond to high levels of specialisation in larval diet. This adds a new dimension to the complexity of functional ecology ^7, 8^. While it has been discussed that functional loss at the parasitoid larval stage has the potential to trigger important cascading effects across ecosystems^19^, a comprehensive understanding of functional risk must account for both larval and adult functional strategies.

### Contrasting patterns of functional distinctiveness and specialisation in larvae *vs* adults

Functional distinctiveness is a form of ecological uncommonness, but it does not necessarily imply ecological specialisation, numerical rarity, or geographic restriction ^3^. A species can be functionally distinct because it exhibits a unique combination of traits, while still being common in abundance or distribution. Conversely, a species may be distinct because it possesses traits that allow it to exploit rare or highly specific resources, effectively making it a functional specialist and a rare species at the same time. This ambiguity complicates the ecological interpretation of the triangular pattern observed in the correlation between larval and adult diet distinctiveness in European bristle flies, where species can be highly distinct in one developmental stage but not in the other. If functionally distinct species are common in space or abundance, the pattern we observed might arise simply from probability: only a minority of species are highly distinct at either stage, so the joint probability of being highly distinct at both stages is inherently low. Alternatively, if distinctiveness reflects ecological specialisation and true rarity, then the absence of species that are highly distinct at both stages may indicate that such a dual specialisation is an unsuccessful evolutionary strategy. Indeed, ecological, numerical, or geographic rarity are known to increase vulnerability to environmental change ^20^, and being twice vulnerable – as a larva and as an adult – may be selectively disadvantageous.

Unlike functional distinctiveness, specialisation always implies that a species has a restricted ecological niche breadth ^21^, being able to exploit a limited set of resources. Here we found an asymmetry in larval *vs* adult diet specialisation, which may arise from spatial constraints imposed by adult foraging niches. In fact, we found that only specialised larvae (those attacking one or two types of hosts) develop into specialised adults. If the adult is specialised in feeding on particular floral resources – which may occur in rare or spatially restricted habitats, specific altitudinal layers, or particular microclimatic conditions – it is likely confined to a narrow spatial domain to efficiently exploit such resources. As a consequence, the larva might be limited by the availability of hosts, which are also confined to the same narrow spatial domain. This shows that selective pressure toward specialisation acts synergistically across developmental stages in some lineages, perhaps because the resources required by both larva (e.g. host) and adult (e.g., flowers) are restricted. This is the case for example for species such as *Sarromyia nubigena* and *Chaetovoria antennalis*, both confined to sparse alpine grasslands and fellfields ^22^ above the tree line, or subalpine species such as *Peleteria popeli* occurring just below the tree line ^23^. Yet, we also found that the opposite is not true, as generalist adults can derive from both generalist and specialised larvae. In other words, it is possible to have a decoupling of functional specialisation from larval to adult stage. The decoupling of diet strategies (as either generalist or specialist) between life stages is not uncommon among entomophagous insects. Some aphidophagous Syrphidae (e.g. *Paragus haemorrhous* and *Eupeodes nielseni*) are known to specialise in the predation of aphids at larval stage, before transforming into adults that feed as generalist palynivores ^24^. Other species of parasitoid flies, such as conopids of the genus *Physocephala*, are characterized by particularly long and narrow snouts as adults, which make them specialised palynivore feeders, but they attack a large number of different hosts as larvae (Sphecidae, bees, bumble bees, and social wasps) which makes them generalist parasites ^25^.

Our classification of larval diet specialisation, in terms of the diversity of defence strategies in the hosts they attack, requires some evolutionary considerations. Bristle flies appear to have colonised lepidopteran hosts relatively late in evolutionary history ^26, 27^, likely diversifying onto an already complex and structured trophic substrate. In this context, caterpillar defence strategies – such as aposematism, gregariousness, concealment, or setosity – most likely evolved as defences against visually oriented predators (such as birds or other insects) or against hymenopteran parasitoids. These traits have then become cues exploited by bristle flies, effectively turning defensive caterpillar phenotypes into parasitoid targets. In other words, these “well-defended” hosts may have represented a less-contested trophic resource for bristle flies, consistent with the concept of “enemy-free space” ^28^. While our analysis was restricted to a subset of easily observable phenotypic defences of caterpillars, other potentially important factors (such as chemical defences or internal physiological barriers) may also influence host vulnerability to parasitism. In fact, it is possible that some of the caterpillars we identified as having no defence traits may have defence mechanisms we did not evaluate here.

### Implications for functional homogenisation in parasitoid insects

The accelerating rates of global environmental change pose a serious threat to functional diversity across ecosystems, favouring a few strategies while negatively impacting others. While the risk of functional loss has historically received much less attention compared to taxonomic loss, growing evidence suggests that the risk of functional homogenisation – the increased similarity in functional trait composition – is of far more concern than the risk of taxonomic homogenisation ^29^, even if the latter is only partly related to the former. Importantly, global environmental change can affect different aspects of functional diversity in a different way. Species with different functional traits respond differently to environmental and anthropogenic pressures, resulting in functional “winners” and “losers” ^30, 31^. Functional homogenisation may arise, for instance, when generalist species replace specialists in response to environmental change (either due to a decline of specialists, an increase of generalists, or both). While specialist species have a competitive advantage over generalists under stable conditions, generalist species tend to be favoured in heterogeneous and perturbed environments ^29^. But the situation is further complicated in parasitoids, as our results show the same species can be either a generalist or a specialist depending on the life stage.

In a previous work ^16^ we found a reshuffling of host specialisation in bristle fly larvae along elevational bands, with species showing specialised larval diet becoming less common at high elevations. This pattern was attributed to the effects of climate change, which resulted in lowland generalist species colonising higher elevations, as already observed in other taxa ^32, 33^. This might depend on generalist feeders being more effective than specialist ones at tracking the upwards expansion of their hosts, or acquiring the ability to exploit novel hosts in newly colonized elevational bands. Our current results imply that the loss of functional diversity might operate in asynchronous ways across larvae and adults. The loss of functionally distinct larvae might reduce the overall capacity to control certain insect hosts, with a consequent risk of herbivory outbreak ^34^. In some cases this also implies the loss of specialised adults, with a consequent reduction in the overall diversity of pollinators within an ecosystem and a risk of decline for certain flowering plant species. While our aim here was to characterise the functional diversity of the European bristle fly fauna, and how it changes across life stages, it is important to clarify that community assemblages play a crucial role in determining which parts of such potential diversity are actually manifested in a given location. In other words, there is both a global and a contextual interpretation of functional diversity ^35^. It is possible, for example, that a species which is globally common from a functional perspective is rare in certain specific contexts. The opposite is also possible, for example functionally distinct strategies can be locally common (e.g. in high-altitude environments).

Global-scale biodiversity monitoring has customarily focussed on taxonomic, sometimes phylogenetic, elements of diversity, while functional diversity has often been underrepresented. Our results show the complex nature of functional diversity in parasitoid insects, with potentially complex risks deriving from functional loss. As parasitoids maintain crucial ecosystem functions across their life stages, understanding how functional homogenisation might affect both larvae and adults across space and time is crucial to anticipate widespread ecosystem consequences.

## Methods

### Bristle fly functional traits

As bristle flies undergo complete metamorphosis and shift their ecological roles between larval and adult stages, we characterized functional traits separately for the two stages. We considered a number of morphological, reproductive, and diet traits that characterise both larvae and adults (Table 1). We attributed to the larval stage all traits connected with the species’ development strategy, including those related to the egg (even if this stage, of course, precedes the larva); we considered the taxonomic order of the attacked host, the attacked stage of the host, the diet breadth, and two features of the egg. For the adults, we considered the shape of the labella, the length of the mouthparts, the overall body length, the shape of the oviscape, and the laying strategies (Dataset S1).

**Table 1.**
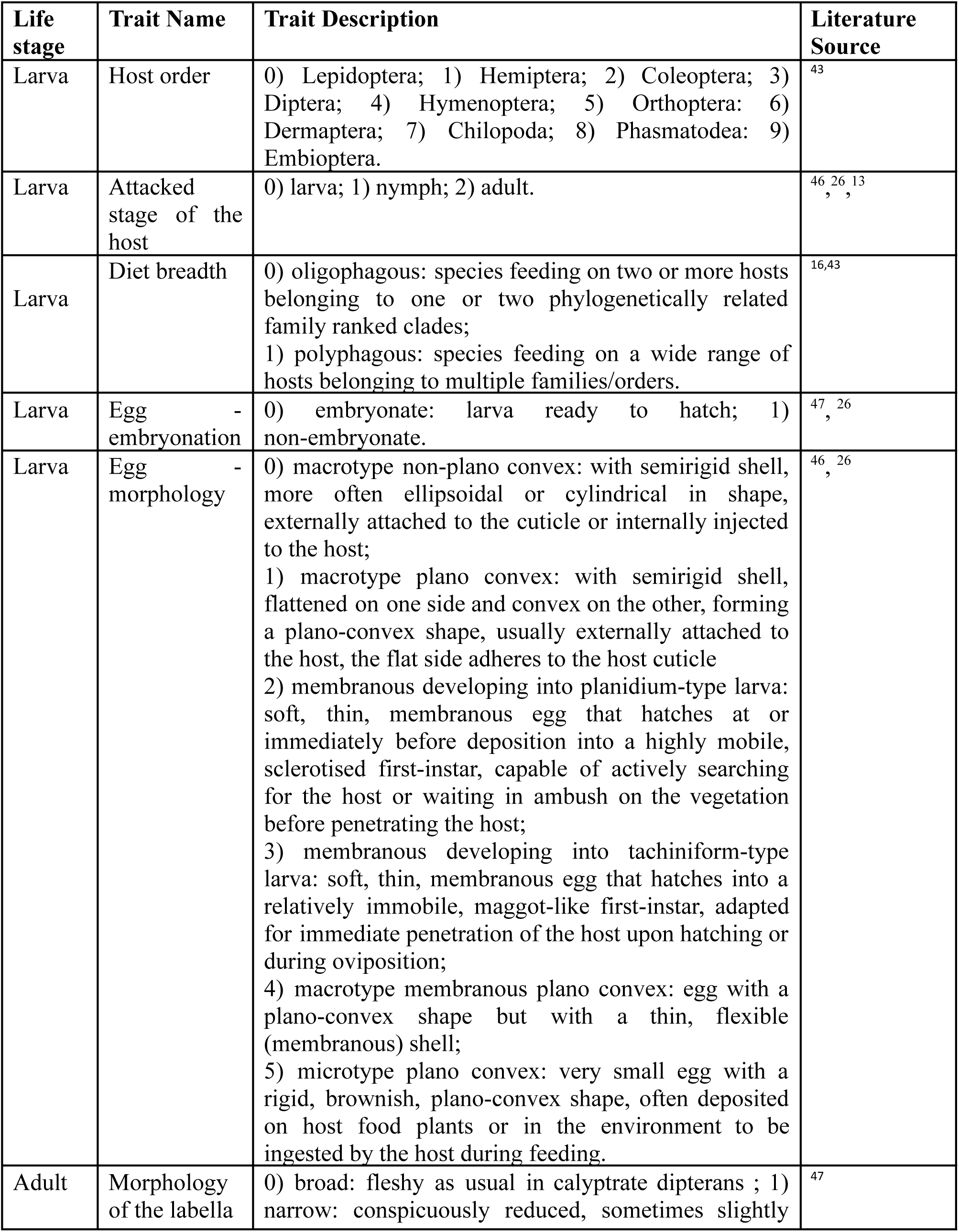
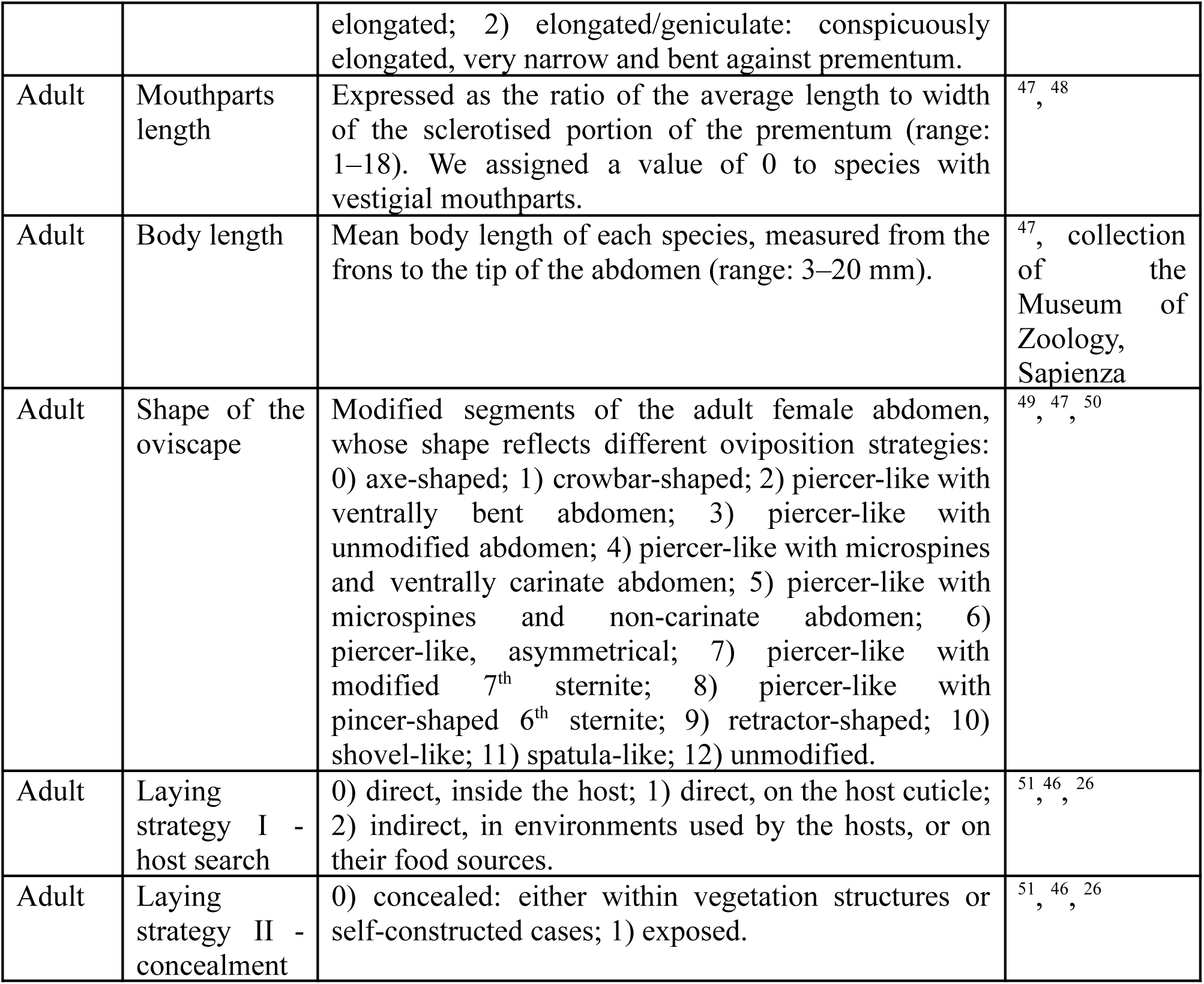
Description of the functional traits of European bristle fly species, at larval and adult stages. Note: we attributed to the larval stage all traits connected with the species’ development strategy, including those related to the egg (even if this stage, of course, precedes the larva).

| Life stage | Trait Name | Trait Description | Literature Source |
| --- | --- | --- | --- |
| Larva | Host order | 0) Lepidoptera; 1) Hemiptera; 2) Coleoptera; 3) Diptera; 4) Hymenoptera; 5) Orthoptera; 6) Dermaptera; 7) Chilopoda; 8) Phasmatodea; 9) Embioptera. | <sup>43</sup> |
| Larva | Attacked stage of the host | 0) larva; 1) nymph; 2) adult. | <sup>46, 26, 13</sup><br>, , |
| Larva | Diet breadth | 0) oligophagous: species feeding on two or more hosts belonging to one or two phylogenetically related family ranked clades;<br>1) polyphagous: species feeding on a wide range of hosts belonging to multiple families/orders. | <sup>16,43</sup> |
| Larva | Egg - embryonation | 0) embryonate: larva ready to hatch; 1) non-embryonate. | <sup>47, 26</sup><br>, |
| Larva | Egg - morphology | 0) macrotype non-plano convex: with semirigid shell, more often ellipsoidal or cylindrical in shape, externally attached to the cuticle or internally injected to the host;<br>1) macrotype plano convex: with semirigid shell, flattened on one side and convex on the other, forming a plano-convex shape, usually externally attached to the host, the flat side adheres to the host cuticle<br>2) membranous developing into planidium-type larva: soft, thin, membranous egg that hatches at or immediately before deposition into a highly mobile, sclerotised first-instar, capable of actively searching for the host or waiting in ambush on the vegetation before penetrating the host;<br>3) membranous developing into tachiniform-type larva: soft, thin, membranous egg that hatches into a relatively immobile, maggot-like first-instar, adapted for immediate penetration of the host upon hatching or during oviposition;<br>4) macrotype membranous plano convex: egg with a plano-convex shape but with a thin, flexible (membranous) shell;<br>5) microtype plano convex: very small egg with a rigid, brownish, plano-convex shape, often deposited on host food plants or in the environment to be ingested by the host during feeding. | <sup>46, 26</sup><br>, |
| Adult | Morphology of the labella | 0) broad: fleshy as usual in calyptrate dipterans ; 1) narrow: conspicuously reduced, sometimes slightly elongated; 2) elongated/geniculate: conspicuously elongated, very narrow and bent against prementum. | <sup>47</sup> |
| Adult | Mouthparts length | Expressed as the ratio of the average length to width of the sclerotised portion of the prementum (range: 1–18). We assigned a value of 0 to species with vestigial mouthparts. | <sup>47</sup>, <sup>48</sup> |
| Adult | Body length | Mean body length of each species, measured from the frons to the tip of the abdomen (range: 3–20 mm). | <sup>47</sup>, collection of the Museum of Zoology, Sapienza |
| Adult | Shape of the oviscap | Modified segments of the adult female abdomen, whose shape reflects different oviposition strategies: 0) axe-shaped; 1) crowbar-shaped; 2) piercer-like with ventrally bent abdomen; 3) piercer-like with unmodified abdomen; 4) piercer-like with microspines and ventrally carinate abdomen; 5) piercer-like with microspines and non-carinate abdomen; 6) piercer-like, asymmetrical; 7) piercer-like with modified 7<sup>th</sup> sternite; 8) piercer-like with pincer-shaped 6<sup>th</sup> sternite; 9) retractor-shaped; 10) shovel-like; 11) spatula-like; 12) unmodified. | <sup>49</sup>, <sup>47</sup>, <sup>50</sup> |
| Adult | Laying strategy I - host search | 0) direct, inside the host; 1) direct, on the host cuticle; 2) indirect, in environments used by the hosts, or on their food sources. | <sup>51</sup>, <sup>46</sup>, <sup>26</sup> |
| Adult | Laying strategy II - concealment | 0) concealed: either within vegetation structures or self-constructed cases; 1) exposed. | <sup>51</sup>, <sup>46</sup>, <sup>26</sup> |

Trait data had missing information for 279 species (Table S1), which we imputed to calculate metrics of functional diversity. Specifically, we imputed missing values for seven categorical variables in our dataset containing morpho-functional traits of bristle flies (Table S1). The imputation was performed using available traits as well as information on species phylogenetic relationships ^36^. We retrieved a phylogenetic tree for bristle flies ^27^ and decomposed the resulting phylogenetic distance matrix into a set of orthogonal eigenvectors using the R package “PVR” ^37^. Phylogenetic information was generally available at the genus level, except for 56 species for which phylogeny was fully resolved at the species level. A total of 79 species instead lacked both trait and phylogenetic data. For these species, we calculated the mean eigenvector values at the genus level and assigned these genus-level means to all congeneric species. We then performed an imputation procedure using the “MissForest” package ^38^. To determine the optimal number of eigenvectors for improving imputation accuracy, we repeated the procedure using datasets containing the first 5, 10, 15, or 20 eigenvectors. To evaluate the effectiveness of the phylogenetically-informed imputation, we computed the Misclassification Error Rate (MER) from the “missForestPredict” package ^39^. This metric is specifically applicable to categorical variables and is calculated as the ratio between the sum of false positives and false negatives, and the sum of total positives and negatives from the confusion matrix. MER values range from 0 to 1, indicating best and worst performance, respectively. The machine learning imputation procedure was generally successful at filling data gaps (MER < 0.2), with 10 eigenvectors as the best-performing set for imputation purposes (Extended Data Fig. 8).

### Analysis of functional distinctiveness

We defined two functional morphospaces for European bristle flies, one for larvae and one for adults, based on their combination of functional traits. Here we aimed to characterise the functional characteristics of individual species with respect to the entire European bristle fly fauna, rather than identifying differences between local communities. Functional spaces were derived using the “funspace” R package ^35^, which estimates the probability of occurrence of each species in a certain portion of the morphospace using kernel density estimation with unconstrained bandwidth selectors. This was achieved by generating a matrix of Gower distances among species, based on both numerical and categorical trait variables, using the R package “vegan” ^40^; we then applied a Constrained Analysis of Principal Coordinates on the distance matrix to generate a larval and an adult morphospaces, using the R package “funrar” ^41^. We mapped each species according to its position in the larval and adult morphospaces. In order to assess the degree of correspondence in functional strategies between larval and adult stages, we extracted the “density” from the position where species occurred in the two morphospaces. We retrieved the species scores from the ordination with the “vegan” R package, and then we extracted the kernel density estimate for each species using the “ks” R package ^42^ for both life stages. This allowed us to evaluate whether species in a “dense” portion of the functional trait space as larvae, also occupied a dense portion as adults.

We then calculated the functional distinctiveness of each species within the functional trait space, with the R package “funrar” ^41^. For this purpose, Gower’s distance matrices were obtained separately for the two life stages of each species. To evaluate whether trait distinctiveness was specifically driven by diet-related traits, we also repeated this analysis by limiting traits to mouthparts length and labella shape for adults, and taxonomic order of the host for larvae. For larvae, the number of different parasitised species, genera, and families was also included. Finally, we compared functional distinctiveness between the two life stages to identify possible patterns of trait-based correspondence. This comparison allowed us to reveal whether the distinctiveness of species’ ecological roles remains steady across metamorphosis, or if levels of differentiation are independent at each life stage.

### Analysis of diet specialisation for lepidopteran feeders

To investigate patterns of ecological specialisation, in addition to distinctiveness, we focused on diet-related traits which could be classified along a generalist-specialist gradient based on expert knowledge. We focused this analysis on lepidopteran feeders only, as these had comprehensive information available on host species which could be retrieved from the literature. To do so, we collected information on caterpillars attacked by bristle flies from Tschorsnig^43^. We preferred to use direct measures of diet specialisation, as opposed to measures derived from analysis of species position in the morphospace, as this approach allows for a more direct interpretation of results along an expert-defined specialisation gradient which reflects Eltonian specialisation ^21^.

We defined larval diet specialisation in terms of their ability to exploit “defence strategies” of the caterpillar hosts. Host defence strategies were assessed using traits associated with the following defensive mechanisms: aposematism, hairiness, gregariousness, and non-exposure (i.e. whether caterpillars forage on exposed *vs* concealed vegetation elements). These traits have previously been identified as ecologically relevant for tachinid parasitism ^19^ due to their role in morphological and behavioural defences. In fact, defence mechanisms of caterpillars (evolved as a defence against predators) can be exploited by bristle fly larvae, as these have a higher chance of hatching success from a host with low predation risk. Each of the 1,194 caterpillar species in our dataset was coded for the binary status (0,1) of the four above-described defensive traits, yielding 13 observed trait combinations out of 16 theoretical ones (i.e. 2^4). Hereafter, we refer to these combinations as “defence strategies”. For each bristle fly species we calculated the number of defence strategies parasitised, in this case a strategy was selected when at least one lepidopteran species with that strategy was parasitised. We assumed that specialists would be able to attack only a limited set of defence strategies, whilst generalists would be able to exploit a wider range of strategies.

We defined adult diet specialisation focussing on the shape of the labella. This trait is a proxy for trophic specialisation ^44^, as the reduction in the number of pseudo-tracheae derived by labella narrowing and elongation is associated with nectar feeding from deep corollas ^45^. Broadly-shaped labella indicate generalist species, narrowly-shaped labella intermediate degrees of specialisation, and elongated/geniculate labella specialist species.

## Acknowledgements

This project was supported by Sapienza University of Rome—Progetti di Ricerca Grandi (CUP B83C23005390005).

## LIST OF EXTENDED DATA FIGURES

**Extended Data Figure 1.**
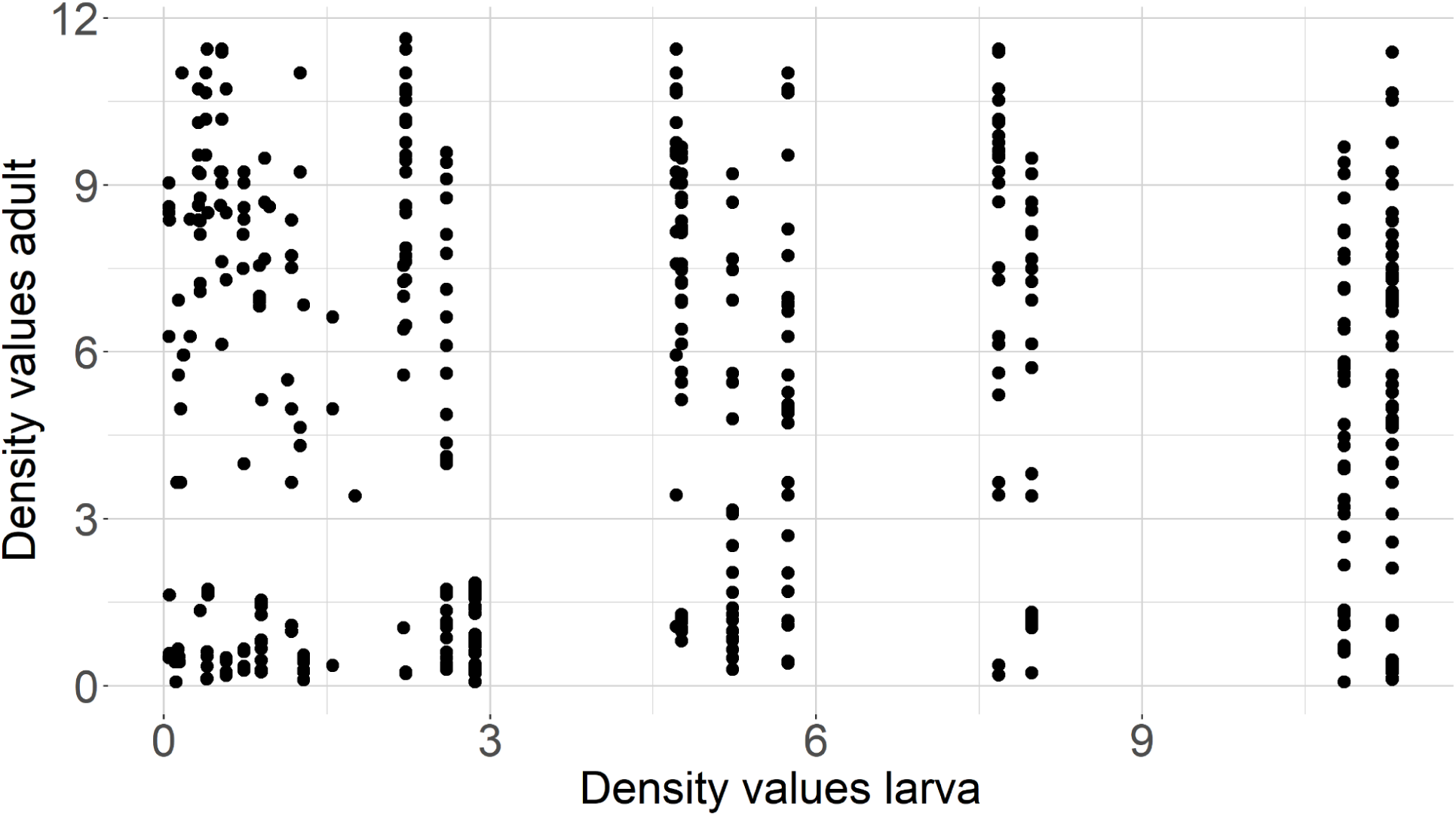
Scatterplot displaying the distribution of density values for each species during the larval and adult life stages, derived from kernel density estimation (KDE) of their positions in the functional morphospace. Each point represents a species, with the x-axis showing the KDE density value in the larval stage and the y-axis showing the KDE density value in the adult stage.

**Extended Data Figure 2.**
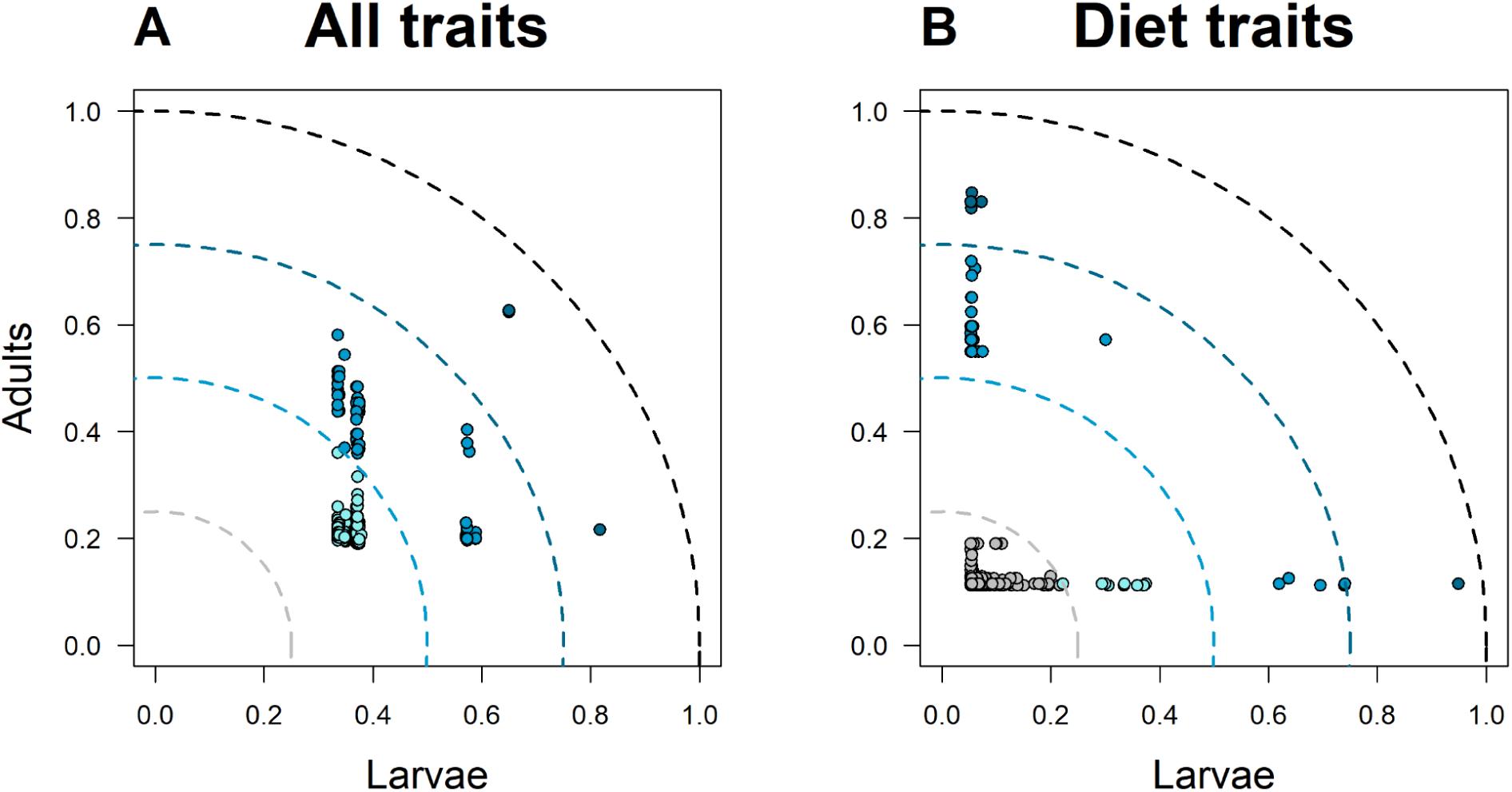
Functional distinctiveness of bristle fly species attacking lepidopteran hosts, at larval (x axis) vs adult (y axis) stage. Panels show results based on (A) all functional traits, and (B) only diet-related traits. The dashed lines represent increased distance from the origin (i.e. increased levels of distinctiveness).

**Extended Data Figure 3.**
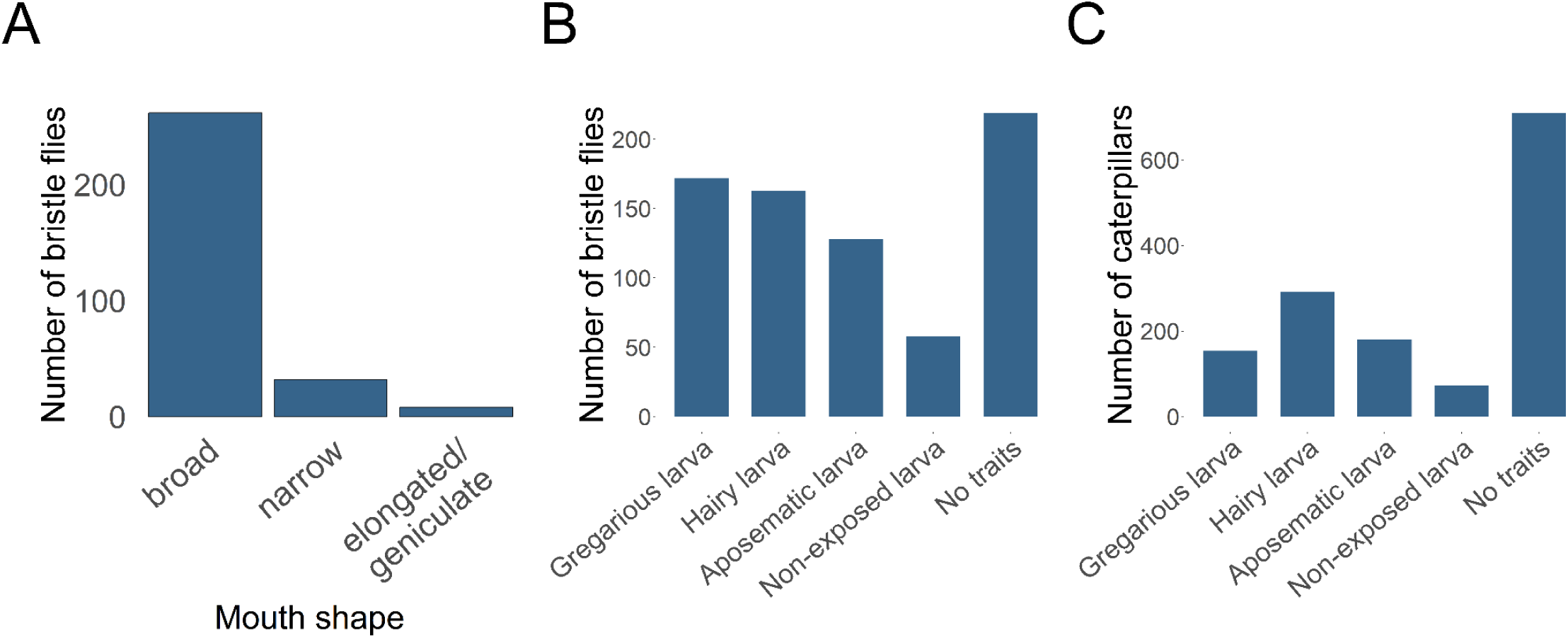
Number of bristle fly species in each diet category based on (A) adult mouthparts (labella shape, used as a proxy of adult diet specialisation) and (B) larval ability to attack caterpillar hosts with certain defence strategies. Panel (C) reports the number of caterpillar species in each defence strategy, for comparison.

**Extended Data Figure 4.**
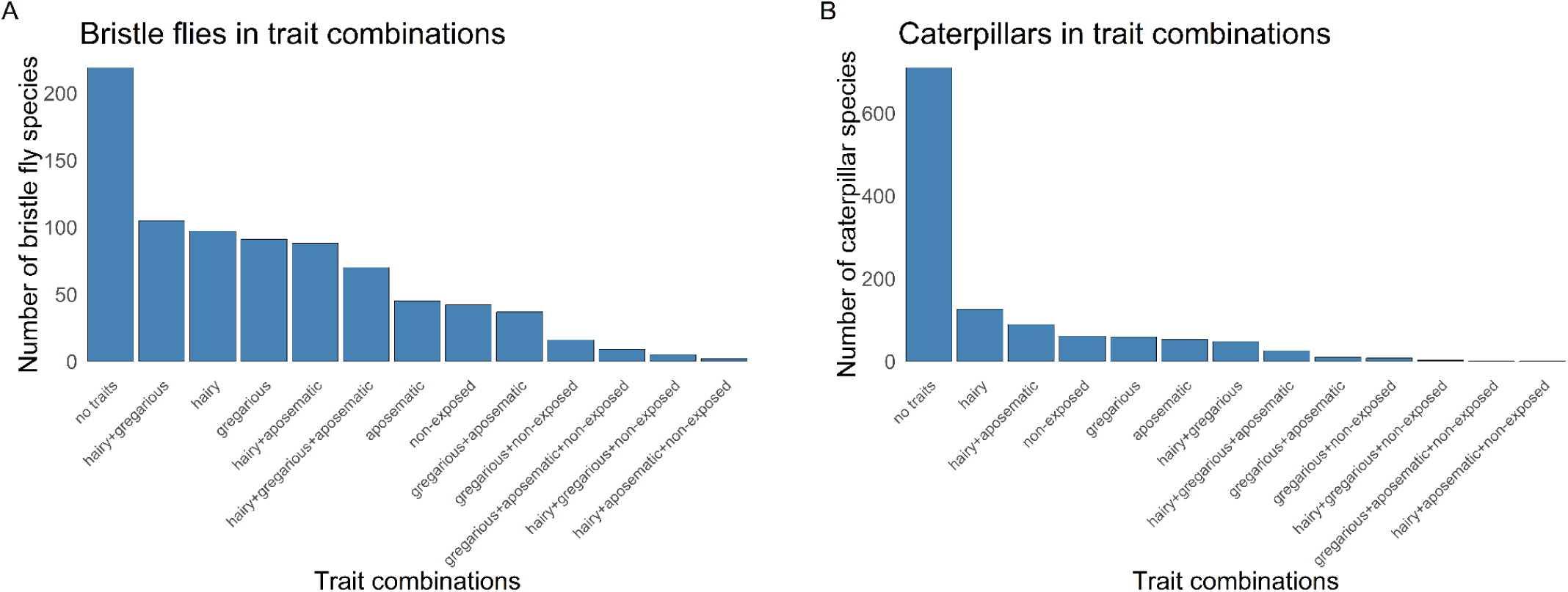
Number of (A) bristle fly species and (B) lepidopteran host species falling within each defence strategy; defence strategies are identified as unique defence trait combinations of the caterpillar hosts.

**Extended Data Figure 5.**
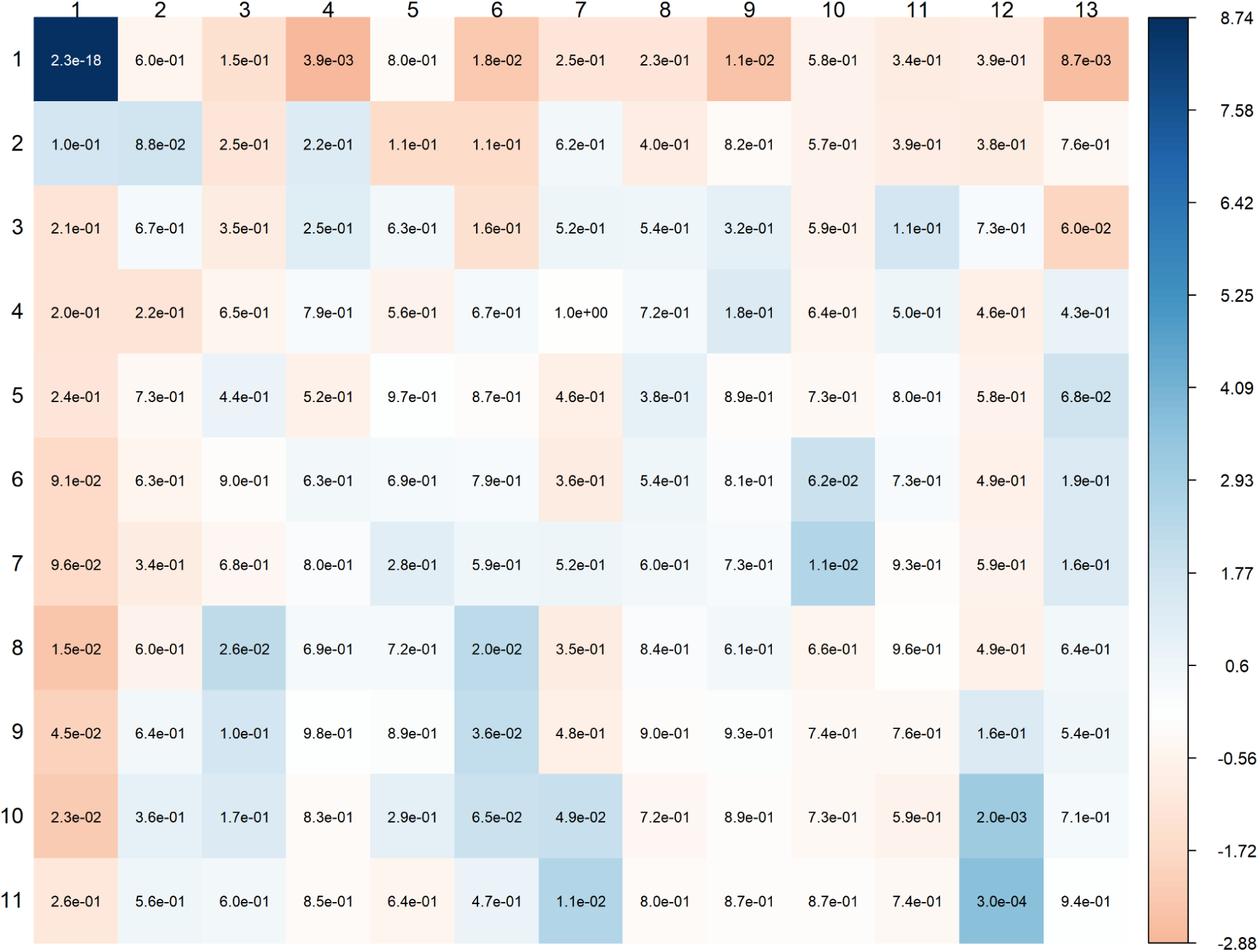
Heatmap displaying the results of a Fisher’s exact test assessing the number of bristle fly species attacking a certain defence strategy (column) for any given number of strategies attacked in total (row). The color intensity of each cell reflects the magnitude of the standardized residuals for each combination of variable levels: high positive values mean there is a higher than expected proportion of species in that combination, very negative values mean the opposite. The numbers inside the cells represent the bilateral p-values associated with the standardized residuals. Defence strategies are coded as follows: 1 “no traits”, 2 “non-exposed”, 3 “aposematic”, 4 “gregarious”, 5 “gregarious + non-exposed”, 6 “gregarious + aposematic”, 7 “gregarious + aposematic + non-exposed”, 8 “hairy”, 9 “hairy + aposematic”, 10 “hairy + aposematic + non-exposed”, 11 “hairy + gregarious”, 12 “hairy + gregarious + non-exposed”, 13 “hairy + gregarious + aposematic”.

**Extended Data Figure 6.**
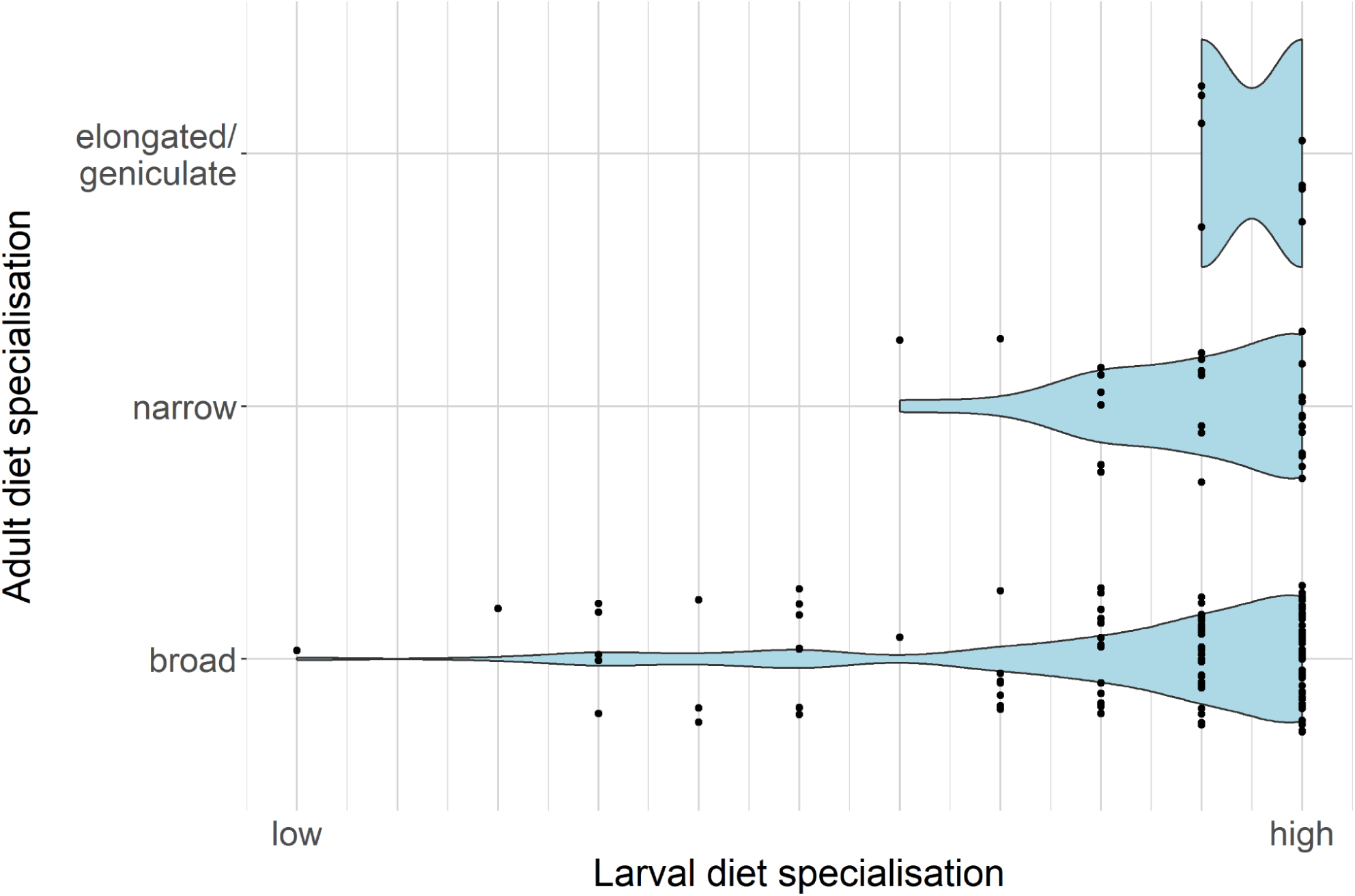
Relationship between diet functional specialisation at larval and adult stages, for species with indirect oviposition. Larval specialisation is represented in terms of the number of host defence strategies attacked by the larva, in decreasing order (i.e. more specialised larvae can only attack a few host types). Adult specialisation is represented in terms of the shape of their mouth part with “broad” being the most generalist form and “elongated” being the most specialised one.

**Extended Data Figure 7.**
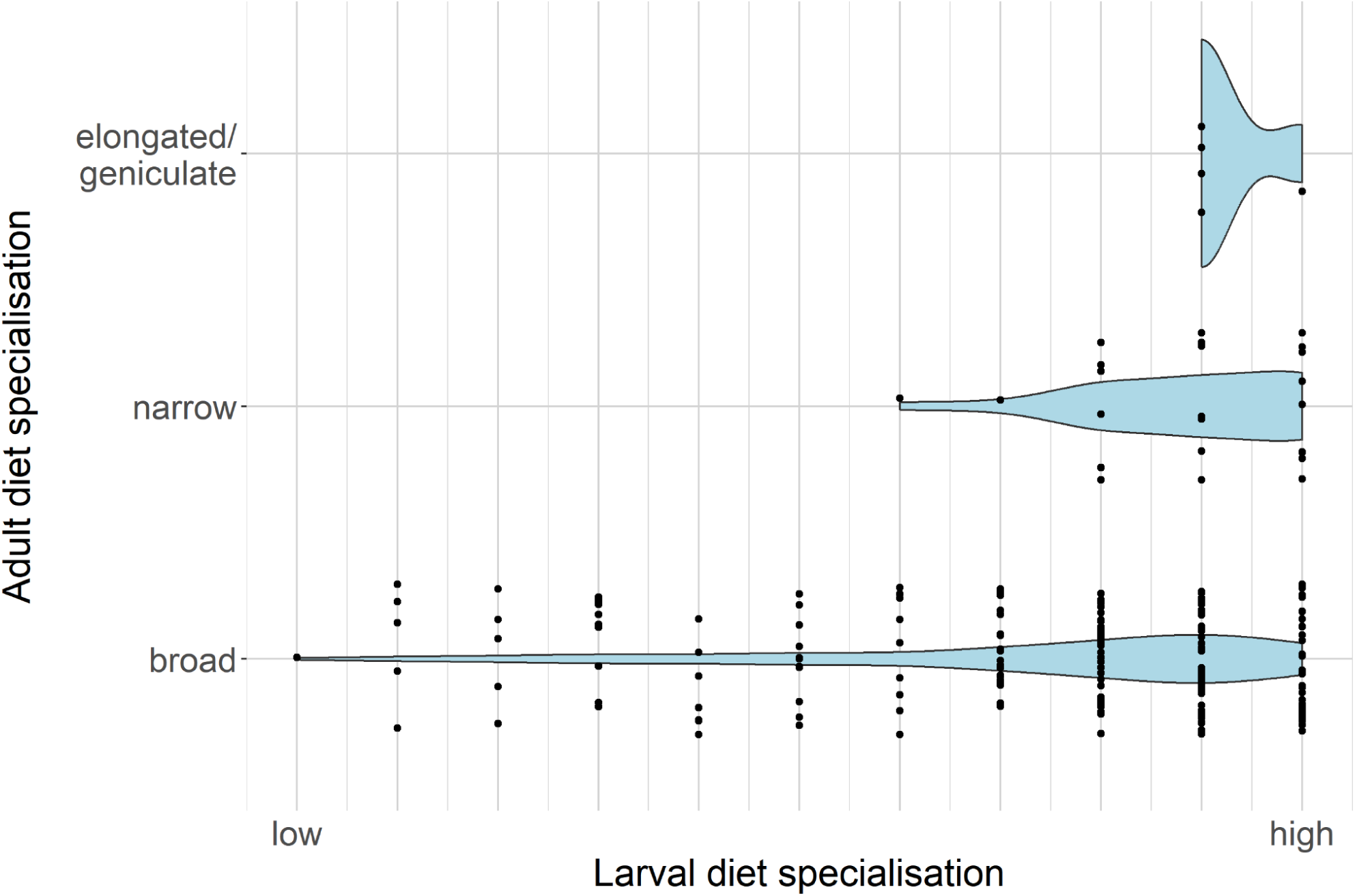
Relationship between diet functional specialisation at larval and adult stages, omitting bristle fly species attacking caterpillars with no defence strategies. Larval specialisation is represented in terms of the number of host defence strategies attacked by the larva, in decreasing order (i.e. more specialised larvae can only attack a few host types). Adult specialisation is represented in terms of the shape of their mouth part with “broad” being the most generalist form and “elongated” being the most specialised one.

**Extended Data Figure 8.**
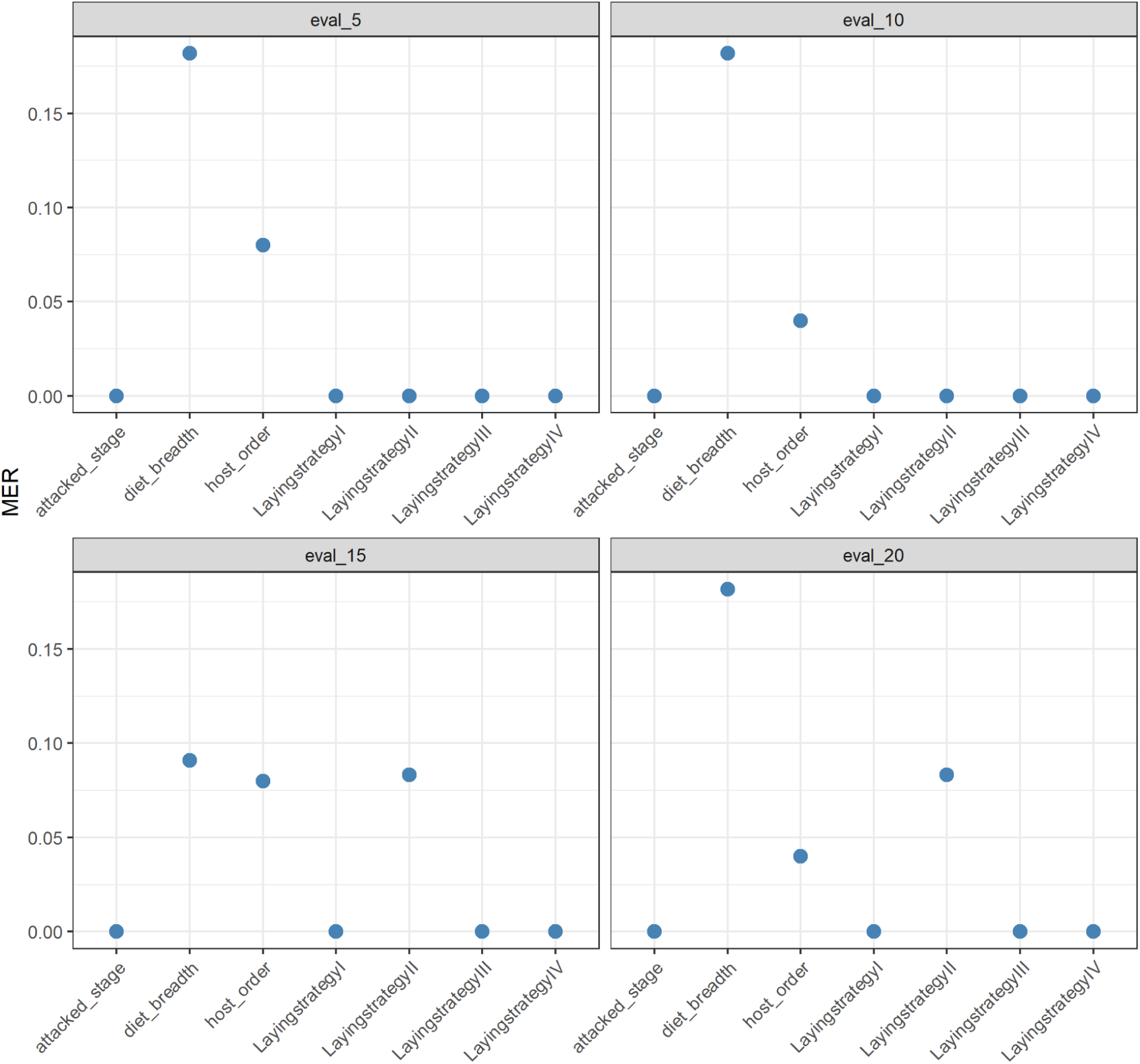
Imputation performance across different levels of phylogenetic eigenvector inclusion. The x-axis reports the imputed columns from the original dataset, whilst the y-axis shows the Misclassification Error Rates (MER), ranging from 0 (best performance) to 1 (worst performance).

## References

1. Petchey, O. L. & Gaston, K. J. Functional diversity: back to basics and looking forward. Ecol. Lett. 9, 741–758 (2006).

2. De Bello, F. et al. Functional trait effects on ecosystem stability: assembling the jigsaw puzzle. Trends Ecol. Evol. 36, 822–836 (2021).

3. Violle, C. et al. Let the concept of trait be functional! Oikos 116, 882–892 (2007).

4. Gagic, V. et al. Functional identity and diversity of animals predict ecosystem functioning better than species-based indices. Proc. R. Soc. B Biol. Sci. 282, 20142620 (2015).

5. Brum, F. T. et al. Global priorities for conservation across multiple dimensions of mammalian diversity. Proc. Natl. Acad. Sci. 114, 7641–7646 (2017).

6. Griffin, J. N. et al. Functionally unique, specialised, and endangered (FUSE) species: towards integrated metrics for the conservation prioritisation toolbox. Preprint at 10.1101/2020.05.09.084871 (2020).

7. Mouillot, D., Villéger, S., Scherer-Lorenzen, M. & Mason, N. W. H. Functional Structure of Biological Communities Predicts Ecosystem Multifunctionality. PLOS ONE 6, e17476 (2011).

8. Leitão, R. P. et al. Rare species contribute disproportionately to the functional structure of species assemblages. Proc. R. Soc. B Biol. Sci. 283, 20160084 (2016).

9. Thomson, L. J., Macfadyen, S. & Hoffmann, A. A. Predicting the effects of climate change on natural enemies of agricultural pests. Biol. Control 52, 296–306 (2010).

10. Van Lenteren, J. C. The state of commercial augmentative biological control: plenty of natural enemies, but a frustrating lack of uptake. BioControl 57, 1–20 (2012).

11. Cingolani, M. F. et al. Dipteran parasitoids as biocontrol agents. BioControl 70, 285–300 (2025).

12. Stireman, J. O., O’Hara, J. E. & Wood, D. M. TACHINIDAE: Evolution, Behavior, and Ecology. Annu. Rev. Entomol. 51, 525–555 (2006).

13. Cerretti, P. & Tschorsnig, H. P. Annotated host catalogue for the Tachinidae (Diptera) of Italy. Stuttg. Beitr. Zur Naturkunde A 3, 305–340 (2010).

14. Lefebvre, V., Villemant, C., Fontaine, C. & Daugeron, C. Altitudinal, temporal and trophic partitioning of flower-visitors in Alpine communities. Sci. Rep. 8, 4706 (2018).

15. Komonen, A., Penttilä, R., Lindgren, M. & Hanski, I. Forest fragmentation truncates a food chain based on an old-growth forest bracket fungus. Oikos 90, 119–126 (2000).

16. Di Marco, M., Santini, L., Corcos, D., Tschorsnig, H. P. & Cerretti, P. Elevational homogenization of mountain parasitoids across six decades. Proc. Natl. Acad. Sci. 120, e2308273120 (2023).

17. O’Hara, J. E. WORLD GENERA OF THE TACHINIDAE (DIPTERA) AND THEIR REGIONAL OCCURRENCE (2012).

18. Pape, T. et al. Fauna Europaea:Diptera–Brachycera. Biodivers. Data J. 3, e4187 (2015).

19. Stireman, J. O. & Singer, M. S. DETERMINANTS OF PARASITOID–HOST ASSOCIATIONS: INSIGHTS FROM A NATURAL TACHINID–LEPIDOPTERAN COMMUNITY. Ecology 84, 296–310 (2003).

20. Sykes, L., Santini, L., Etard, A. & Newbold, T. Effects of rarity form on species’ responses to land use. Conserv. Biol. 34, 688–696 (2020).

21. Devictor, V. et al. Defining and measuring ecological specialization. J. Appl. Ecol. 47, 15–25 (2010).

22. Ziegler, J. & Tschorsnig, H. P. An overview of all the recorded species in the study area and in South Tyrol, with new data from recent years. Stud. Dipterol. 21, 312–406 (2016).

23. Tschorsnig, H. P., Ziegler, J. & Herting, B. Tachinid flies (Diptera: Tachinidae) from the Hautes-Alpes, France. Stuttg. Beitr. Zur Naturkunde Ser. Biol. 656, (2003).

24. Speight, M. C. D. SPECIES ACCOUNTS OF EUROPEAN SYRPHIDAE (DIPTERA). Syrph the Net, the database of European Syrphidae. Syrph the Net publications. 65, (2024).

25. Stuke, J.-H. Taxonomic notes on Western Palaearctic Conopidae (Diptera). Zootaxa 4178, (2016).

26. Cerretti, P. et al. Signal through the noise? Phylogeny of the Tachinidae (Diptera) as inferred from morphological evidence. Syst. Entomol. 39, 335–353 (2014).

27. Stireman, J. O., Cerretti, P., O’Hara, J. E., Blaschke, J. D. & Moulton, J. K. Molecular phylogeny and evolution of world Tachinidae (Diptera). Mol. Phylogenet. Evol. 139, 106358 (2019).

28. Jeffries, M. J. & Lawton, J. H. Enemy free space and the structure of ecological communities. Biol. J. Linn. Soc. 23, 269–286 (1984).

29. Clavel, J., Julliard, R. & Devictor, V. Worldwide decline of specialist species: toward a global functional homogenization? Front. Ecol. Environ. 9, 222–228 (2011).

30. Dornelas, M. et al. A balance of winners and losers in the Anthropocene. Ecol. Lett. 22, 847–854 (2019).

31. Leung, C., Rescan, M., Grulois, D. & Chevin, L.-M. Reduced phenotypic plasticity evolves in less predictable environments. Ecol. Lett. 23, 1664–1672 (2020).

32. Moritz, C. et al. Impact of a century of climate change on small-mammal communities in Yosemite National Park, USA. Science 322, 261–4 (2008).

33. Rödder, D., Schmitt, T., Gros, P., Ulrich, W. & Habel, J. C. Climate change drives mountain butterflies towards the summits. Sci. Rep. 11, 1–12 (2021).

34. Brodmann, P. A., Wilcox, C. V. & Harrison, S. Mobile Parasitoids may Restrict the Spatial Spread of an Insect Outbreak. J. Anim. Ecol. 66, 65 (1997).

35. Carmona, C. P., Pavanetto, N. & Puglielli, G. funspace: An R package to build, analyse and plot functional trait spaces. Divers. Distrib. 30, 1–14 (2024).

36. Penone, C. et al. Imputation of missing data in life-history traits datasets: which approach performs the best? Methods Ecol. Evol. 5, 961–970 (2014).

37. Diniz-Filho, J. A. F., de Sant’Ana, C. E. R. & Bini, L. M. An Eigenvector Method for Estimating Phylogenetic Inertia. Evolution 52, 1247–1262 (1998).

38. Stekhoven, D. J. & Bühlmann, P. MissForest--non-parametric missing value imputation for mixed-type data. Bioinforma. Oxf. Engl. 28, 112–8 (2012).

39. Albu, E., Gao, S., Wynants, L. & Calster, B. V. missForestPredict -- Missing data imputation for prediction settings. Preprint at 10.48550/arXiv.2407.03379 (2024).

40. Oksanen, J., et al. vegan: Community Ecology Package. The R Foundation 10.32614/cran.package.vegan (2001).

41. Grenié, M., Denelle, P., Tucker, C. M., Munoz, F. & Violle, C. funrar: An R package to characterize functional rarity. Divers. Distrib. 23, 1365–1371 (2017).

42. Duong, T. ks: Kernel Density Estimation and Kernel Discriminant Analysis for Multivariate Data in R. J. Stat. Softw. 21, 1–16 (2007).

43. Tschorsnig, H. P. Preliminary host catalogue of Palaearctic Tachinidae (Diptera). 1–480 (2017).

44. Gilbert, F. & Jervis, M. Functional, evolutionary and ecological aspects of feeding-related mouthpart specializations in parasitoid flies. Biol. J. Linn. Soc. 63, 495–535 (1998).

45. Krenn, H. W. Insect Mouthparts: Form, Function, Development and Performance. Springer Nature, (2019).

46. Tschorsnig, H. P. & Herting, B. The tachinids (Diptera: Tachinidae) of central Europe: identification keys for the species and data on distribution and ecology. 150 (1994).

47. Cerretti, P. I Tachinidi Della Fauna Italiana (Diptera Tachinidae), Con Chiave Interattiva Dei Generi Ovest-Paleartici. Vol. I (Cierre Edizioni, 2010).

48. Cerretti, P., Tschorsnig, H. P., Lopresti, M. & Di Giovanni, F. MOSCHweb — a matrix-based interactive key to the genera of the Palaearctic Tachinidae (Insecta, Diptera). ZooKeys 205, 5–18 (2012).

49. Herting, B. Das weibliche Postabdomen der calyptraten Fliegen (Diptera) und sein Merkmalswert für die Systematik der gruppe. Zeitschrift für Morphologie und Ökologie der Tiere. 45, 429–46 (1957).

50. Tschorsnig, H. P. & Richter, V. A. Contributions to a Manual of Palaearctic Diptera (with special reference to flies of economic importance). Science Herald, Budapest Vol. 3: Higher Brachycera, 691–827 (1998).

51. Herting, B. Biologie der westpalaarktischen Raupenfliegen, Dipt., Tachinidae. Monographien zur angewandte Entomologie 16, 1–188 (1960).

