## Supplementary material for "Two forms, two functions: functional strategies of parasitoid bristle flies and their larvae": Table S1

***Table S1*** *List of the traits with missing data*

| **Trait name** | **Percentage of missing data species** |
|---|---|
| Taxonomic order of the host | 11.99 |
| Attacked stage of the host | 11.08 |
| Diet breadth | 33.63 |
| First feature of the egg | 1.04 |
| Second feature of the egg | 1.04 |
| Laying strategy I | 3.12 |
| Laying strategy II | 22.29 |

**Dataset S1** Functional traits of European bristle flies at larval and adult stage, as described in Table 1. Provided as a separate file.
